# Inertial coupling between swing and stance legs shapes ankle torque profile during running

**DOI:** 10.64898/2026.09.17.752346

**Authors:** Jinsung Jung, Hyerim Lim, Sukyung Park

## Abstract

The spring-loaded inverted pendulum (SLIP) model has effectively explained center of mass (CoM) and ground reaction force (GRF) dynamics during human gait despite its simplicity, describing both walking and running within the same mechanical principles. A recent study extended the explanatory capability of the SLIP model to joint dynamics by reproducing ankle joint dynamics during walking. However, during running, the model did not capture the empirical ankle torque profile, suggesting that such mechanical unification may not hold at the joint level. Because the previous model represented the body as a single point mass, it neglected leg inertia, which may become important during running as rapid motion generates substantial angular momentum. To address this limitation, we propose an extended two-mass model that separates the whole-body mass into the swing-leg and the remaining body mass, thereby explicitly incorporating swing-leg inertia into stance-leg joint dynamics. The swing leg is modeled using a linear spring and a torsional spring at the hip joint for its radial and rotational motion. Through inertial coupling, swing-leg dynamics influence the stance-leg ankle torque. Consequently, the proposed model reproduces the symmetric, single-peaked ankle torque profile observed experimentally while preserving the CoM and GRF of the previous model. These findings suggest that the discrepancy of the previous model during running may be explained by the omission of swing-leg inertia. By accounting for the gait-mode-dependent contribution of swing-leg inertia, the proposed model provides a unified mechanical framework for extending simple gait models from CoM-level behavior to joint dynamics.

## 1. Introduction

The most fundamental task of human gait is to move the whole-body mass forward through interaction with the ground. Accordingly, understanding the relationship between center of mass (CoM) motion, which represents whole-body behavior, and the ground reaction force (GRF) is central to understanding gait dynamics. In particular, using a simple spring-loaded inverted pendulum (SLIP) model composed of a lumped mass and spring, Geyer et al. demonstrated that two distinct gait modes, walking and running, can be explained under the same principles of passive dynamics (Geyer et al., 2006). It has also been reported that GRF propulsion and the corresponding CoM dynamics across different gait speeds can be reproduced by varying spring stiffness, and the same framework remains applicable across different age groups and walking conditions, including load carriage (Kim and Park, 2011; Hong et al., 2013; Lee et al., 2014). However, despite its ability to reproduce whole-body-level behavior, the virtual leg inherently cannot capture joint-level information because it does not explicitly include joints.

To address this limitation, Lim and Park sought to explain lower-limb joint-level dynamics by introducing structural modifications to the passive dynamics of the SLIP model (Lim and Park, 2018). Their model successfully reproduced major characteristics of lower-limb dynamics during walking, particularly ankle dynamics, which play a major role in gait (Neptune et al., 2001; Farris and Sawicki, 2012). However, when the same approach was applied to running, the model produced an asymmetric ankle torque profile that differed from experimental observations. This discrepancy suggests that the extension of SLIP-based mechanical framework from CoM-level to joint-level dynamics remained incomplete.

One possible reason for this discrepancy during running is the simplification of the body as a single lumped mass. During walking, lower-limb motion is relatively slow, and neglecting inertial effects associated with limb motion was a reasonable assumption. In contrast, during running, lower limbs move more rapidly and over larger ranges of motion (Novacheck, 1998). Consequently, the swing leg generates substantial angular momentum during the stance phase, and swing-leg inertia may influence stance-leg joint dynamics.

In previous studies, the swing leg has primarily been interpreted in terms of its kinematic role, for example as a controller contributing to stability or establishing appropriate landing conditions (Seyfarth et al., 2003; Blum et al., 2010; Rashty et al., 2014). However, relatively few studies have examined the influence of swing-leg inertia on stance-leg joint dynamics during running. Therefore, examining the influence of swing-leg inertia may help explain why the conventional single-mass model does not sufficiently reproduce ankle torque during running.

In this study, we propose an extended-SLIP model that separates the swing-leg mass from the remaining body mass, thereby explicitly incorporating the effect of swing-leg inertia on stance-leg joint dynamics. To represent the radial and rotational motions of the swing leg, the swing-leg mass is connected to the remaining body mass through a linear spring and a torsional spring at the hip joint, respectively. This configuration allows swing-leg inertia to influence the ankle torque of the stance leg through inertial coupling. The model was validated by comparing the simulated ankle torque during the stance phase of running with experimental data. This study examines whether incorporating swing-leg inertia can extend the explanatory capability of a simple spring-mass framework from CoM-level behavior to joint-level dynamics during running.

## 2. Methods

### 2.1. Proposed extended-SLIP model description

To incorporate the inertial effects of swing-leg on stance-leg joint dynamics during running, we extended model of Lim and Park by separating swing-leg mass from the remaining body mass (Fig. 1).

#### 2.1.1. Model configuration and assumption

In the previous model, a point mass *M*, representing the whole body, was supported by a stance-leg spring *k* compliantly connected to an off-centered curvy foot. The stance-leg spring *k* consisted of two springs connected in series: *k_leg_* above the ankle and *k_foot_* below the ankle. Because *k_leg_* and *k_foot_* were connected at a massless joint, *k* was defined as their equivalent stiffness. The curvature of the curvy foot was represented by *R*, and its degree of off-centering was represented by *d*. To account for swing-leg inertia, the single lumped mass *M* in the previous model was separated into the swing-leg mass *m_sw_* and the remaining body mass *m*. The values of *m_sw_* (∼16% of *M*) and *m* (∼84% of *M*) were determined based on anatomical mass distribution (Dempster, 1955; Clauser et al., 1969), and the swing-leg mass ratio *μ_sw_* was defined as follows:

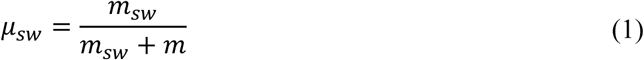

During running, the swing-leg length varies substantially, and the swing leg rapidly rotates in forward direction. Therefore, *m* and *m_sw_* were connected by a linear spring *m_sw_*, while a torsional spring *m_sw_* connected *m_sw_* to a virtual torso oriented vertically with respect to the ground. The two linear springs were assumed to be connected through a pivot joint at *m*. Therefore, the torque generated by the torsional spring does not act directly on the stance leg. Instead, reaction force generated by torsional spring is transmitted to the stance leg through the shared pivot, thereby influencing stance-leg dynamics.

#### 2.1.2. State variables and equations of motion

The state variables of the model shown in Fig. 1 were defined as follows. For the stance leg, the length and angle of spring *k* were denoted by *l* and *θ*, respectively. The angle *θ* was defined as positive in the clockwise direction with respect to the downward vertical. For the swing leg, the length and angle of spring *k_sw_* were denoted by *l_sw_*, and *φ*, respectively. The angle *φ* was defined as positive in the clockwise direction with respect to the upward vertical. The rest angle of the torsional spring *φ*_0_ was set to -170°, corresponding to a natural swing leg orientation close to the standing posture (Fig. 1a).

**Figure 1.**
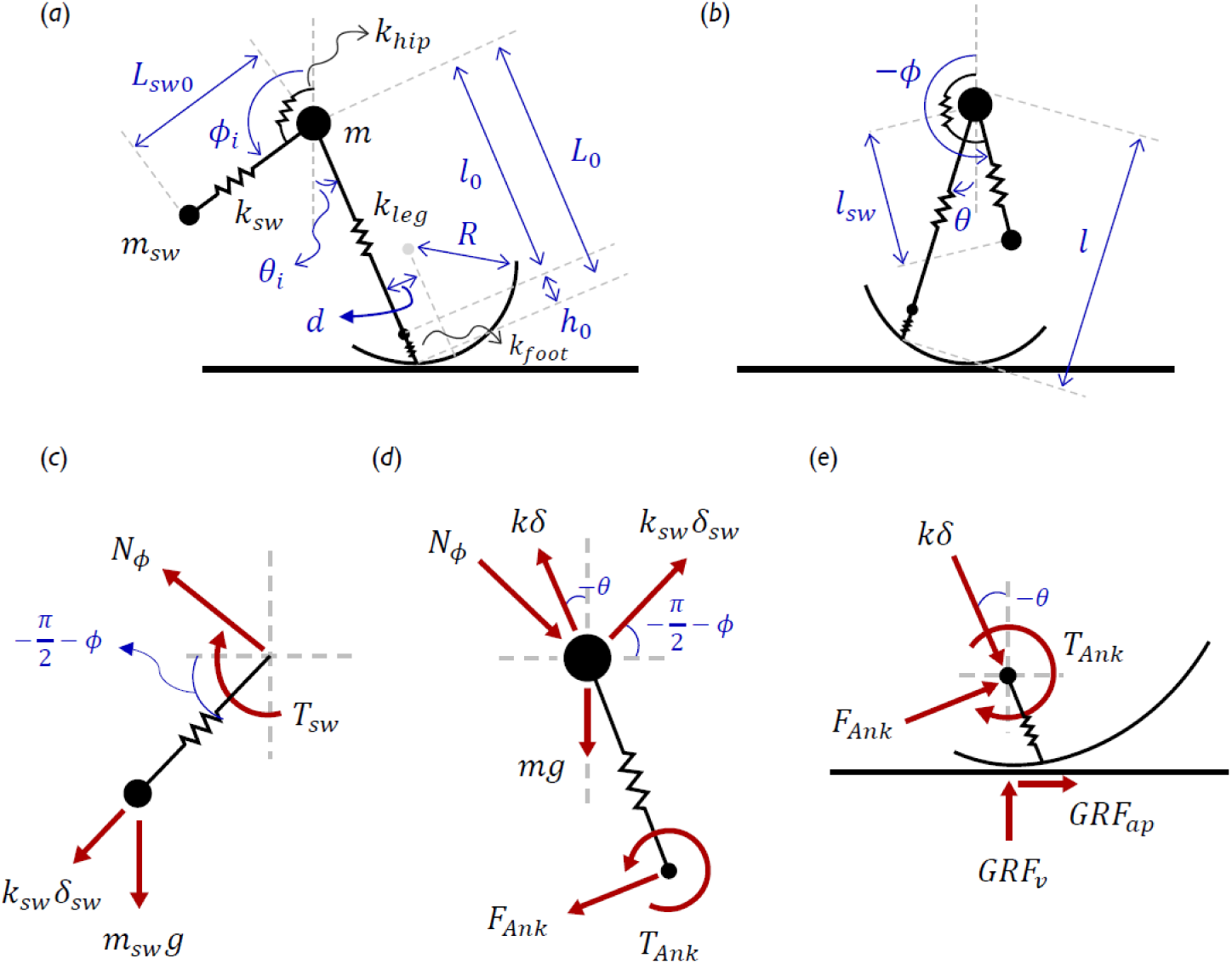
Schematic of the proposed model including swing-leg mass. (*a*) Model configuration and the state variables, parameters, and initial conditions at heel strike, (*b*) state variables of the model at an arbitrary time point during the stance phase, free-body diagrams of (*c*) the swing leg mass, (*d*) the remaining body excluding the swing leg, and (*e*) the massless foot.

Based on the rest lengths of the stance and swing legs, *L*_0_ and *L_sw0_*, respectively, the stance- and swing-leg spring compressions were defined as *δ* = *L*_0_ − *l* and *δ_sw_* = *L_s_*_w0_ − *l_sw_*, respectively. Because the present analysis was limited to the stance phase of running, the equations of motion for stance phase were derived using a Newtonian method as follows.

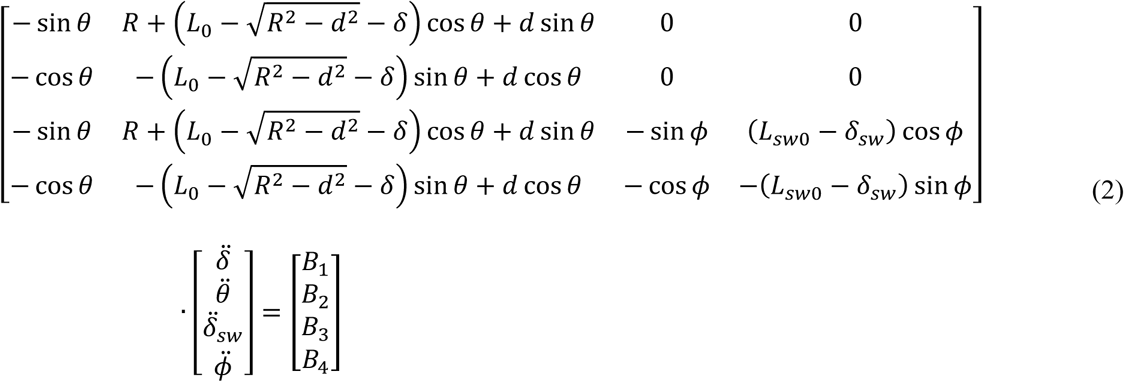

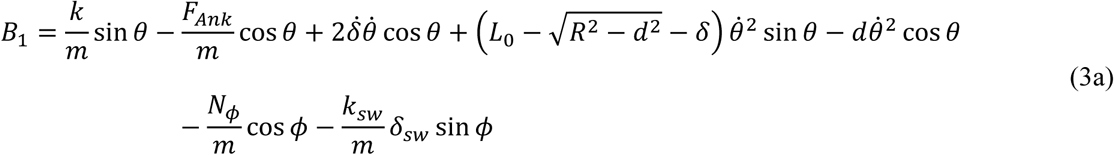

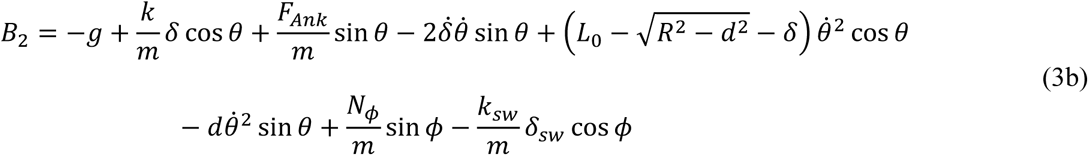

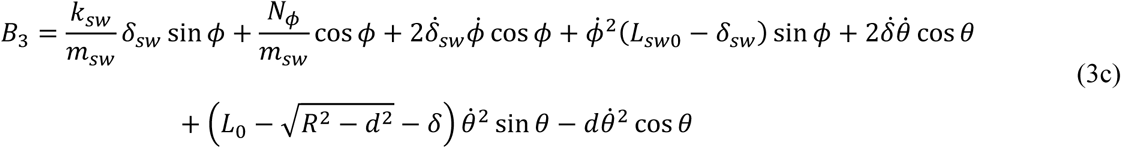

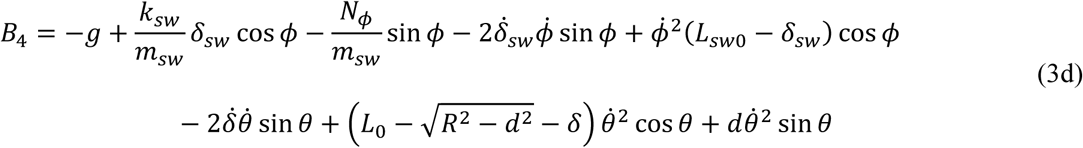

In the equations of motion, *N_φ_* and *F_Ank_* denote the reaction force generated by the torsional spring and the constraint force required to maintain stance-leg alignment on the off-centered curvy foot, respectively. These forces were calculated as follows:

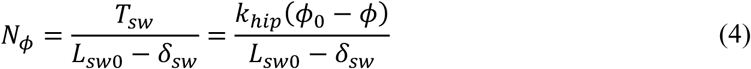

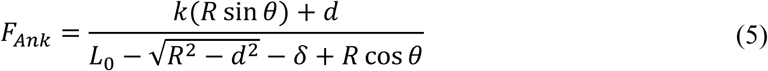

where *T_sw_* denotes the torque generated by the swing-leg torsional spring.

#### 2.1.3. Parameter setting and model simulation

The stiffness *k* and initial leg angle *θ_i_* were selected to best reproduce the experimental data at the prescribed running speed, as in previous SLIP studies. The stance-leg rest length *L*_0_ was set slightly longer than the typical SLIP-model value of 1 m because it represents the distance to the CoM of the remaining body mass rather than the whole-body CoM. Parameters introduced to represent the foot structure in the previous model (Lim and Park, 2018), including *R*, *d*, *r* (=ℎ_0_/*l*_0_), and the stiffness ratio *ρ* (=*k_foot_*/*k_leg_*), were maintained at the same values as in the previous model.

For the newly introduced swing leg, the linear spring stiffness *k_sw_* was initially determined from *k*/*m* = *k_sw_* /*m_sw_* to approximately match the natural frequencies of the stance and swing legs. Because their motions were mechanically coupled, *k_sw_* was adjusted around this estimate to best reproduce the experimental behavior. The rest length *L_sw0_* and the initial angle *φ_i_* were set based on their experimental values at the initial state. The torsional spring stiffness *k*_ℎ*ip*_ and rest angle *φ*_0_ were determined based on experimental hip joint torque and angle data during the running swing phase.

Initial conditions inherited from the previous model were retained, whereas the newly introduced initial conditions were set close to the experimental values.

**Table 1.** Model parameters and initial conditions used for simulation.

| Model parameters of previous model |  |  |  |  |  |  |  |  |
| --- | --- | --- | --- | --- | --- | --- | --- | --- |
| $m$ | $\mu_{sw}$ | $k$ | $L_0$ | $R$ | $d$ | $r$ | $\rho$ | $\theta_i$ |
| 60.5 kg | 0.161 | 17 kN/m | 1.1 m | 0.3 m | 0.11 m | 0.2199 | 1.8265 | -13.5 ° |
| Additional model parameters of proposed model |  |  |  |  |  |  |  |  |
| $k_{sw}$ | $L_{sw0}$ | $\dot{\delta}_{sw,i}$ | $k_{hip}$ | $\dot{\phi}_i$ | $\phi_0$ | $\phi_i$ | | |
| 4.9 kN/m | 0.5 m | 1 m/s | 3.86 Nm/° | -1.77 rad/s | -170 ° | -160 ° |  |  |

Model simulations were performed in MATLAB R2022a (MathWorks, USA), and the nonlinear differential equations were numerically solved using the ode45 function.

#### 2.1.4. Model outputs: CoM and ankle dynamics

For model validation, we examined whether incorporating swing-leg inertia improved stance-leg ankle dynamics while preserving the CoM dynamics successfully reproduced by the conventional lumped-mass model. To this end, ankle torque was evaluated together with the CoM trajectory and GRF calculated from the simulated model data, and the results were compared with experimental data.

Because the proposed model consists of two point masses, the whole-body CoM trajectory was calculated as the mass-weighted average of their trajectories. The anterior-posterior (a-p) and vertical positions of *m* and *m_sw_*, denoted by (*x*, *y*) and (*x_sw_*, *y_sw_*), respectively, were calculated using Eqs. (6) and (7). The position of the CoM, (*x_CoM_*, *y_CoM_*), was then calculated using Eq. (8).

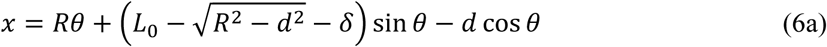

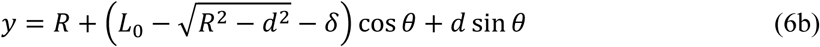

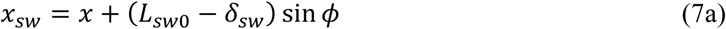

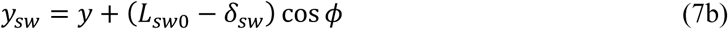

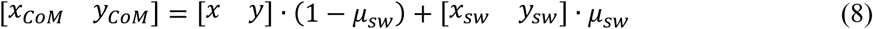

The a-p and vertical components of the GRF, denoted by *GRF_ap_* and *GRF_v_*, respectively, were calculated as the sum of the stance-spring force and the ankle constraint force. The ankle torque was defined as the corresponding constraint torque, as follows:

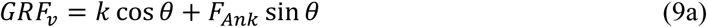

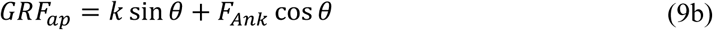

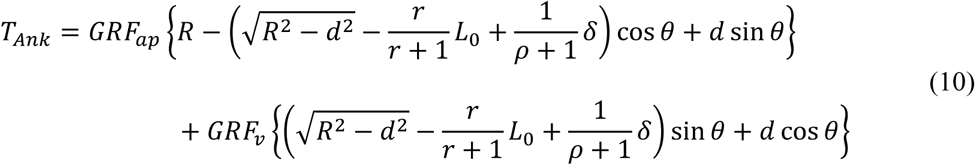

### 2.2. Experiment

#### 2.2.1. Participants and protocol

To compare the proposed model outputs with experimental data, eight healthy adult males with no history of lower-limb musculoskeletal disorders participated in the experiment (age: 24.3 ± 4.4 years; height: 174.8 ± 5.8 cm; mass: 69.3 ± 8.8 kg). Walking data were additionally collected to compare the swing-leg inertia effect between walking and running. All experimental procedures were approved by the Institutional Review Board (KH2023-250), and written informed consent was obtained from all participants prior to the experiment. All methods were performed in accordance with relevant ethical guidelines and regulations. Each participant performed walking (1.3 m/s) and running (3.4 m/s) at fixed speeds on an instrumented treadmill for 8 minutes. A total of 300 steady-state stances, each defined from heel strike to toe-off of one foot, were used for analysis.

#### 2.2.2. Data acquisition and processing

Whole-body kinematics were measured at 100 Hz using a 13-camera motion capture system (MX-T50, VICON, UK) with 40 reflective markers placed on anatomical landmarks. GRFs were measured at 1000 Hz using two force plates embedded in the treadmill (FP6012, Bertec, USA). All data were filtered using a zero-phase 4^th^-order Butterworth low-pass filter with a cutoff frequency of 10 Hz. Lower-limb joint dynamics were calculated from synchronized motion capture and GRF data using a seven-segment lower-body model in Visual3D v6 (HAS-Motion, Canada). Heel strike and toe-off in each stride were identified using a vertical GRF threshold of 8% of body weight. CoM motion was calculated by double integration of the GRF measured from the force plates (Donelan et al., 2002). The angular momentum of the swing leg about the CoM, 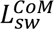, was calculated as follows:

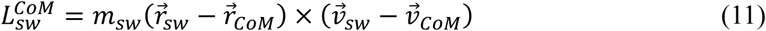

where 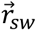 and 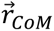 denote the position vectors of swing-leg CoM and whole-body CoM respectively, and their velocity vectors are denoted as 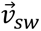 and 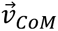, respectively.

### 2.3. Model evaluation and analysis

The effect of incorporating swing-leg inertia was evaluated primarily from the ankle torque, while the CoM trajectory and GRF were examined to determine whether the proposed model preserved the whole-body dynamics reproduced by the previous model. The experimental GRF was normalized by each participant’s body weight, whereas swing-leg angular momentum and ankle torque were normalized by body mass. Participant-averaged profiles were then obtained separately for each variable and compared with the model outputs. The overall profiles were evaluated qualitatively, while the timings of peak ankle torque and vertical GRF during the stance phase were compared quantitatively. CoM and ankle displacements were also compared as supplementary measures.

Additionally, we compared the accumulation pattern of joint dynamics throughout the stance phase using the cumulative ankle torque impulse, *J_T_*_A*nk*_, normalized by the total ankle torque impulse over the stance phase. We also compared the propulsion-to-braking ratio (P-B ratio), defined as the ratio of torque impulses during the propulsion to that during the braking period. The two periods were separated at 50% of the stance phase (Shorter and Rouse, 2020). The two metrics were defined as follows.

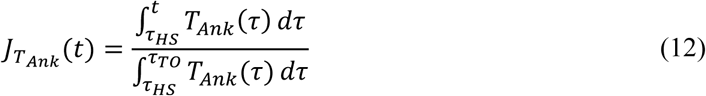

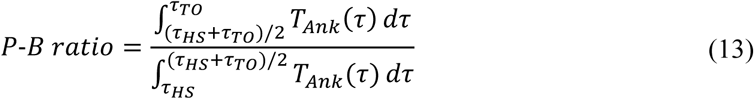

In Eqs. (12) and (13), *τ_HS_* and *τ_T_*_0_ denote heel-strike and toe-off times, respectively, and *T_Ank_*(*τ*) denotes the ankle joint torque. Moreover, we examined the effect of *μ_sw_* on ankle torque reproduction by varying it from 0.05 to 0.35 while maintaining the total body mass. For each *μ_sw_*, the ankle torque profile and P-B ratio were compared with the experimental data.

## 3. Results

The proposed model incorporating swing-leg inertia reproduced stance-leg ankle joint dynamics during running more closely than the previous model without swing-leg mass (Lim and Park, 2018) (Fig. 2). The experimental ankle torque exhibited a relatively symmetric profile, with a peak at approximately 54% of the stance phase. In contrast, the previous model produced an asymmetric profile biased toward the latter half of stance, with a peak at approximately 64% of the stance phase. However, the proposed model incorporating swing-leg inertia reproduced a more symmetric profile with a peak at 53% of the stance phase, closely matching the experimental data (Table 2).

**Figure 2.**
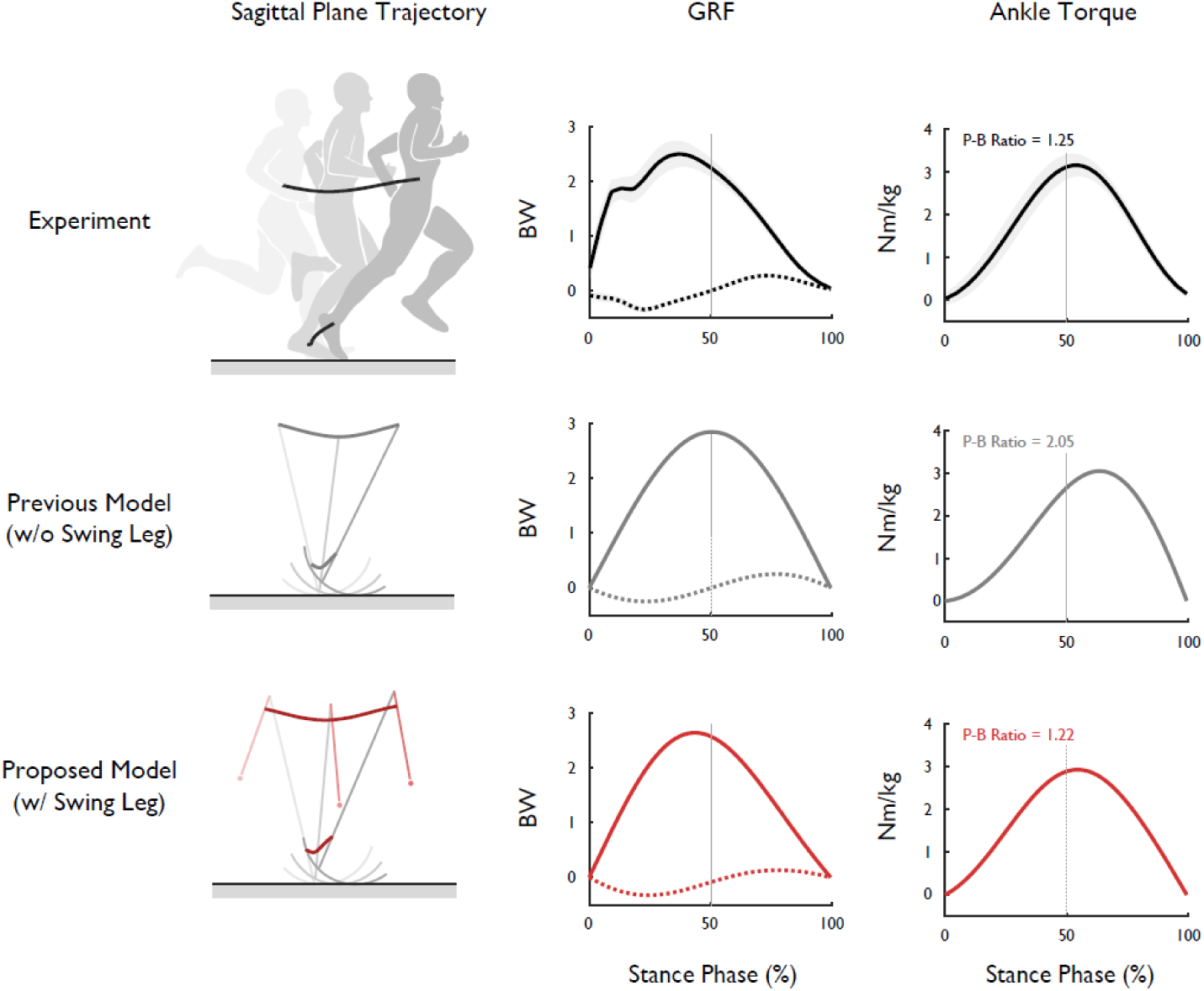
CoM and ankle dynamics results at a running speed of 3.4 m/s for the experiment, the previous model without a swing leg, and the proposed model with a swing leg, shown from top to bottom. The trajectory plots are shown in the sagittal plane, with the horizontal axis representing the direction of progression and the vertical axis representing the vertical direction. From left to right, the columns show the trajectories of the CoM and ankle joint, the vertical (solid line) and a-p (dashed line) GRFs, and the mass-normalized ankle torques. The P-B ratio, representing the degree of late-stance bias in the ankle torque curve, is shown in the upper-left corner of each ankle torque plot. The vertical dashed line in the GRF and ankle torque plots indicates 50% of the stance phase. shaded area around the experimental curve represents ± standard deviation. (*P-B ratio: Experiment 1.25, Previous model 2.05, Proposed model 1.22)

Minor but consistent improvements were also observed in the reproduction of whole-body dynamics, including the GRF and sagittal-plane CoM trajectory. Compared with the previous model, the proposed model generated a vertical GRF profile that was slightly more biased toward the early stance phase and a wider CoM trajectory. It also produced a greater vertical ankle displacement, more closely matching the experimental data (Fig. 2, first and second columns; Table 2).

The improvement in ankle torque reproduction was also observed in the P-B ratio. The previous model showed a P-B ratio of 2.05, substantially higher than the experimental value of 1.25, indicating that the ankle contribution during the latter propulsion period was overestimated relative to that during the early braking period. In contrast, the proposed model reproduced a P-B ratio of 1.22, close to the experimental value.

**Table 2.** CoM and ankle dynamics metrics experimentally observed and reproduced by the previous and proposed models.

|  | Metric | Previous | Proposed | Experiment |
| --- | --- | --- | --- | --- |
| Ankle dynamics | Ankle torque peak timing (%) | 64 | 53 | 54 |
|  | P-B ratio | 2.05 | 1.22 | 1.25 |
|  | Vertical ankle trajectory (cm) | 6.0 | 7.5 | 11.9 |
| CoM dynamics | Vertical GRF peak timing (%) | 51 | 43 | 37 |
|  | A-P CoM trajectory (m) | 0.69 | 0.77 | 0.79 |

The previous model showed a different pattern of increase in *J_T_*_A*nk*_ throughout the stance phase, whereas the proposed model closely reproduced the experimental pattern (Fig. 3). Although the experimental data and both models exhibited S-shaped *J_T_*_A*nk*_ curves, the previous model showed a delayed increase during the early stance phase, followed by a more concentrated increase during the latter half of stance.

**Figure 3.**
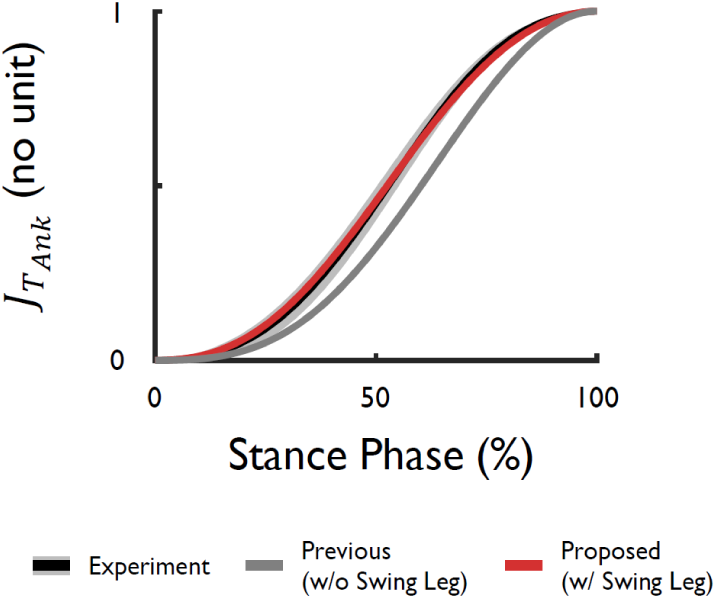
Phase-normalized cumulative ankle torque impulse *J_T_*_A*nk*_ during the running stance phase. Black indicates the experimental result, gray indicates the previous model, and red indicates the proposed model. The shaded area around the experimental curve represents ± standard deviation.

The swing-leg mass ratio *μ_sw_* substantially affected the ankle torque profile in the proposed model, with *μ_sw_* around 0.15-0.2 producing profiles most similar to the experimentally observed ankle torque (Fig. 4a). At low *μ_sw_*, the torque profile resembled that of the previous model, exhibiting a single peak during the latter half of the stance phase. However, excessively high *μ_sw_* (e.g., 0.35) resulted in a double-peaked profile that deviated substantially from the experimental observation.

**Figure 4.**
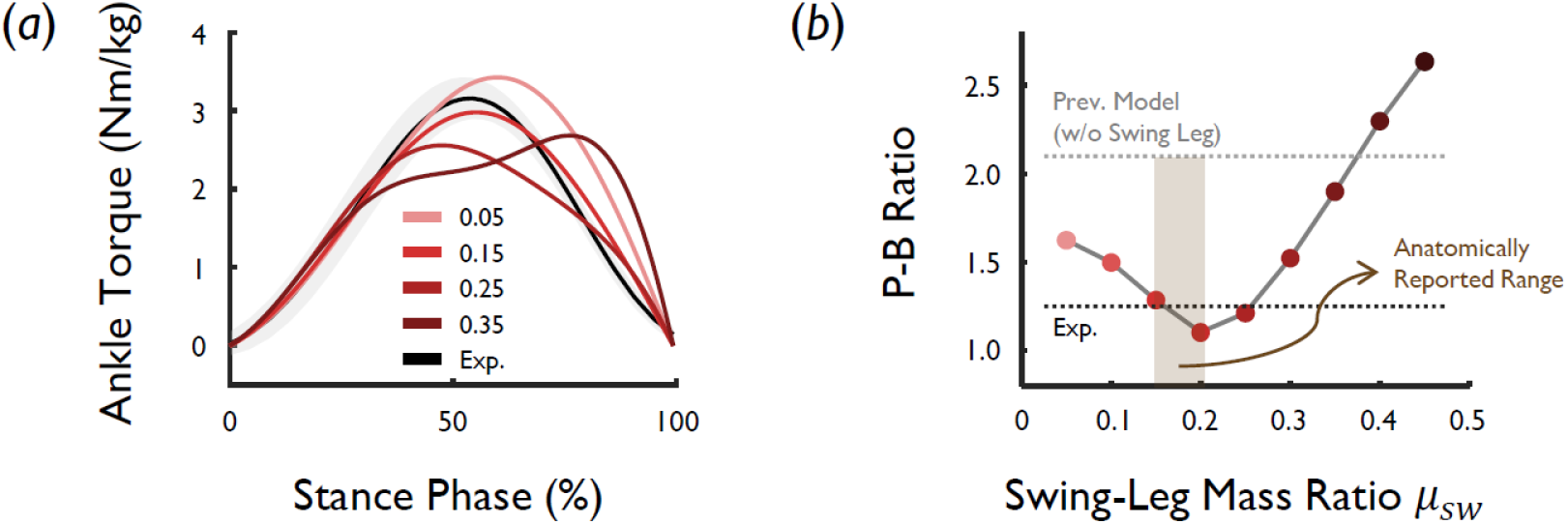
Effects of swing-leg mass ratio *μ_sw_* on ankle torque reproduced by the proposed model. (*a*) Ankle torque profiles at different swing-leg *μ_sw_*. The solid black line indicates the experimentally observed value, and the shaded areas represents ± standard deviation. The red lines indicate the reproduced model results, with darker lines corresponding to higher *μ_sw_*. (*b*) P-B ratio as a function of swing-leg ratio. The light gray horizontal dashed line indicates the P-B ratio of the previous model, while the dark gray horizontal dashed line indicates the experimental P-B ratio. The light brown shaded region indicates the anatomically reported range (Dempster, 1955; Clauser et al., 1969).

These changes were also reflected in the P-B ratio (Fig. 4b). As *μ_sw_* increased from 0.05 to 0.2, the P-B ratio approached the experimental value. At *μ_sw_* greater than 0.2, the P-B ratio increased again and progressively diverged from the experimental value. In particular, within the anatomically reported range of single-leg mass ratios (approximately 0.15-0.2; Dempster, 1955; Clauser et al., 1969), the P-B ratio closely matched the experimental value.

The experimentally observed angular momentum of the swing leg about the CoM during the stance phase was substantially greater during running than during walking (Fig. 5a). During walking, the angular momentum of the swing leg about the CoM (hereafter, swing-leg angular momentum) remained small, with a mean magnitude below 0.03 m²/s. In contrast, during running, it generally ranged from 0.05 to 0.1 m²/s and reached values approximately three times as large as those observed during walking.

The previous model did not incorporate a separate swing leg and therefore could not reproduce this angular momentum. In the proposed model, the simulated angular momentum generally remained within a range similar to that observed experimentally. Its peak occurred at 40% of the stance phase, close to the experimental peak at 38%. However, the simulated angular momentum decreased more rapidly during late stance than the experimental angular momentum.

**Figure 5.**
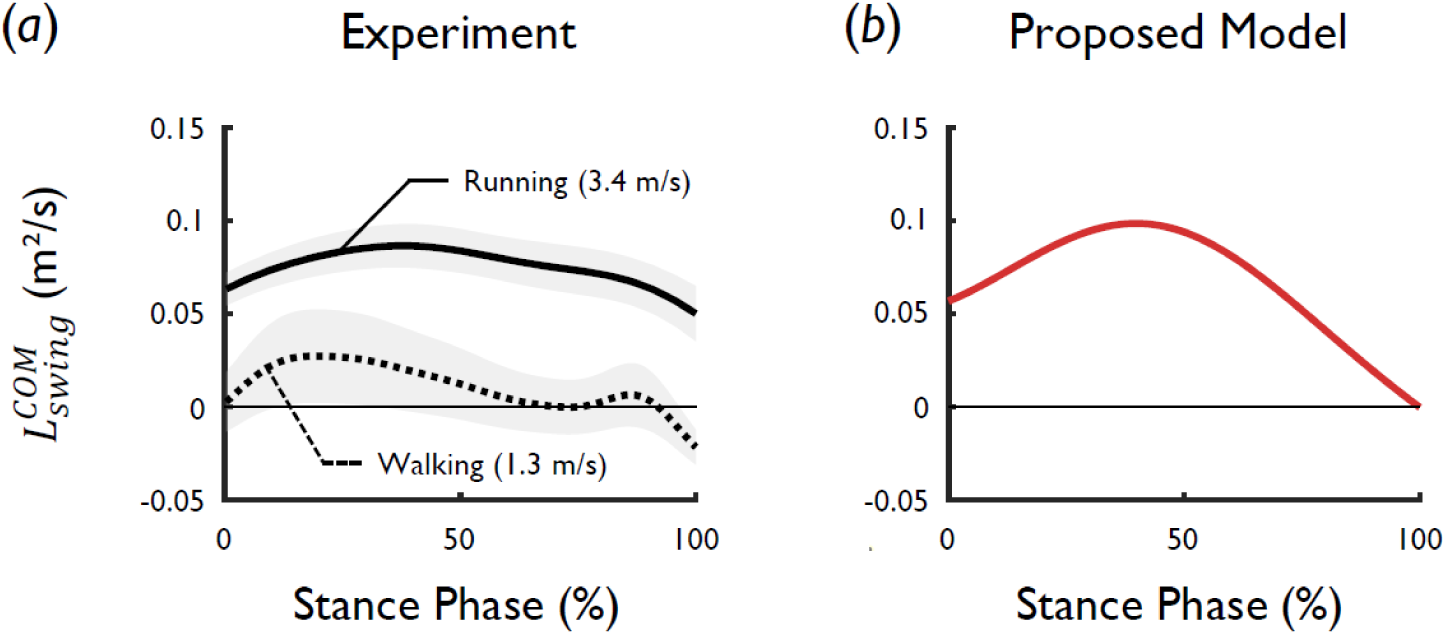
Mass-normalized angular momentum of the swing leg about the CoM during the stance phase. (*a*) Experimentally observed results. The solid and dashed lines indicate running (3.4 m/s) and walking (1.3 m/s), respectively. Shaded areas represent ± standard deviation. (*b*) Results reproduced by the proposed model. The thin horizontal black line in each plot indicates the zero reference.

## 4. Discussion

The results of this study demonstrate that incorporating swing-leg inertia is important for reproducing joint-level dynamics during running. The proposed model substantially reduced the late-stance bias in the ankle torque profile while preserving the major characteristics of the CoM and ankle trajectories and GRFs reproduced by the previous model (Fig. 2).

During running, segmental movements become larger (Novacheck, 1998), and the contribution of swing-leg inertia and angular momentum exchange between the legs increases, resulting in substantial angular momentum of the swing leg about the CoM (Fig. 5a). Therefore, the assumptions that were reasonable for walking, a single mass and negligible swing-leg inertia, no longer hold at the case of running. By explicitly accounting for the swing-leg inertia, the proposed model generates angular momentum of the swing-leg mass about the CoM with a magnitude and peak timing comparable to those observed experimentally (Fig. 5b).

The motion of the incorporated swing-leg mass generated additional inertial loading at the shared pivot. This loading changed the mechanical demand on the stance leg and consequently reshaped the ankle torque profile. As a result, the late-stance bias observed under the single-mass assumption is resolved, and the ankle torque is reconstructed into a more symmetric profile that more closely resembles the human. From the perspective of system angular momentum, the swing leg may thus act as an angular-momentum-bearing component that redistributes mechanical effects between the swing and stance legs.

The present results provide insight into how the mechanical unification between the two gait modes may be established at the joint level. Geyer et al. showed that CoM-level dynamics during walking and running can be explained using the same model and argued that the difference between the two gait modes lies primarily in the energy level (Geyer et al., 2006). However, when Lim and Park extended this framework to the joint level, joint dynamics were successfully reproduced only during walking, which appeared to conflict with this unified paradigm (Lim and Park, 2018). The present findings suggest that extending the unified CoM-level paradigm proposed by Geyer et al. to the joint-level dynamics may require consideration of gait-mode-dependent mechanical effects, particularly swing-leg coupling during running. Under this condition, this study shows that the unified paradigm of walking and running can be extended to joint-level dynamics.

Previous studies modeled swing-leg dynamics only for swing phase during walking (Song et al., 2016; Choi et al., 2025), implemented the swing leg to describe step-to-step transitions (Lim and Park, 2019), or examined the role of swing-leg motion in transition stability (Seyfarth et al., 2003; Blum et al., 2010; Rashty et al., 2014). In contrast, the present study demonstrates that the swing leg, acting as a dynamic mass, can affect the stance-leg ankle torque profile through inertial coupling. Thus, the swing leg may contribute not only to step transition and stability as previously recognized, but also to stance-leg joint dynamics.

The proposed model most closely reproduced the experimental ankle torque within the anatomically reported range of single-leg mass ratios (0.15-0.2) (Dempster, 1955; Clauser et al., 1969) (Fig. 4). When the mass ratio *μ_sw_* was excessively small, the angular momentum of the swing leg diminished, and the ankle torque profile approached that of the previous single-mass model (Lim and Park, 2018). Conversely, excessively high *μ_sw_* produced an unrealistic double-peaked torque profile. These results support that the proposed model extension was based on biomechanically meaningful representation of swing-leg mass rather than arbitrary parameter adjustment.

Despite these implications, the proposed model has several limitations. First, the model focused only on the stance phase, and the flight phase and phase-to-phase transitions were not implemented. Therefore, a complete limit cycle was not evaluated. Future work should integrate the present stance-phase model with an appropriate flight-phase formulation. Second, because the swing leg was simplified as a lumped point mass, the model could not describe detailed internal joint dynamics of the swing leg or their specific effects on stance-leg dynamics. Third, this present analysis was limited to a single running speed. Because running speed may alter swing-leg angular momentum as well as the corresponding model parameters and initial conditions, future studies should evaluate the proposed model over a broader range of running speeds. The small swing-leg angular momentum observed experimentally during walking (Fig. 5a) suggests that swing-leg inertia would have only a minor influence under walking conditions. Therefore, the proposed model may approach the behavior of the previous model during walking and preserve its ability to reproduce walking dynamics (Lim and Park, 2018). Future studies should directly evaluate the proposed model during walking to verify this expectation and further assess the unified joint-level paradigm discussed above.

In summary, this study demonstrated that explicitly incorporating swing-leg inertial coupling improved the reproduction of stance-leg ankle torque during the running stance phase while preserving the major characteristics of CoM-level dynamics. These findings provide a mechanistic pathway for extending the unified gait mode paradigm proposed by Geyer et al. from the CoM level (Geyer et al., 2006) to joint-level dynamics, and may inform simplified modeling approaches for applications such as assistive device design and motion planning.

## CRediT authorship contribution statement

**Jinsung Jung:** Writing – original draft, Writing – review & editing, Conceptualization, Methodology, Software, Validation, Formal analysis, Investigation, Data curation, Visualization. **Hyerim Lim:** Validation, Writing – review & editing, Analysis. **Sukyung Park:** Writing – review & editing, Resources, Supervision.

## Ethics statement

All participants provided written informed consent, and the experimental procedures were approved by the KAIST Institutional Review Board (KH2023-250).

## Declaration of competing interest

The authors declare that they have no known competing financial interests or personal relationships that could have appeared to influence the work reported in this paper.

## Funding

This work was supported by the National Research Foundation of Korea (NRF) grant funded by the Korea Government (MSIT) (No. RS-2024-00356657).

## Notes

### Competing Interest Statement

The authors have declared no competing interest.

